# Rapid estimation of cryptic adult abundance and breeding dynamics in a critically endangered elasmobranch from close-kin mark recapture

**DOI:** 10.1101/2022.02.24.481858

**Authors:** TA Patterson, R Hillary, P Feutry, R Gunasakera, J Marthick, RD Pillans

## Abstract

Elasmobranchs are one of the most highly-threatened vertebrate taxa. Estimating abundance of spawning adults is often extremely challenging, yet crucial for prioritization of conservation measures. Emblematic of these challenges, the speartooth shark (*Glyphis glyphis*, Müller and Henle, 1839) was initially known only from rare specimens collected in a few tropical regions river systems of Australia and Papua New Guinea. Listed as critically endangered in Australia (where only six adults have ever been recorded) and endangered by the IUCN, such scarcity and limited distribution prevent direct assessments of abundance or reproductive biology. We used close-kin mark-recapture to estimate the abundance of mature *G. glyphis* in a genetically isolated population in the Wenlock River, Queensland, Australia. From 224 juvenile/sub-adults and 2 adult samples taken over three years (2013-2016), 46 half-sibling and 33 full-sibling pairs were found. The adult population was estimated at 897 (80% Credible interval 531 – 1684) with a sex ratio (based on mitochondrial DNA) highly skewed to males (females – 0.09: males – 0.91). Based on juveniles sampled in different years that shared a mother, 71% of females were estimated to skip-spawn. In an average year we estimate 44 breeding females occupy the system. Importantly, these methods constitute a viable and relatively rapid approach to obtain robust estimates of absolute abundance and other key population parameters for similar rare species.

## Introduction

A fundamental quantity to monitor in conservation settings is the abundance of breeding adults. Yet, in some species, breeding aged individuals can be cryptic, even if juveniles may be encountered. This study considers a member of the genus Glyphis, the river sharks, which are found the western Indo-Pacific (Li et al. 2015), Papua New Guniea (White et al. 2015) and Northern Australia (Thorburn and Morgan 2004; Compagno, White, and Last 2008; Stevens, Pillans, and Salini 2005; Pillans et al. 2009). For over a century the three species in this genus were known only from type specimens collected in the 19th century and were generally considered extinct (Li et al. 2015). Following discovery of Australian species, *G. glyphis* and *G. garriki* in the late 20th century, the species were conservation listed under IUCN listing and in Australia under the Environmental Protection of Biodiversity and Convseration (EPBC) act of 1999, due to their apparently limited range and rarity. In Australia and Papua New Guinea only 6 adult *G. glyphis* individuals have ever been recorded (Pillans, unpublished data; White et al. 2015).

If species as secretive and poorly understood as river sharks are subject to anthropogenic influence, their populations may be put at risk before their status is even established (Haque and Das 2019). When data collection rates are very slow, conventional survey methods such as visual surveys, mark-release-recapture are often infeasible or simply too slow to establish baseline abundance. In marine systems, non-target species are often monitored only via trends in bycatch data from commercial fisheries(e.g. Jordaan, Santos, and Groeneveld 2018). The problems of characterising non-target populations from fisheries data are well documented. These difficulties are further exacerbated in the case of rare, long-lived and low productivity species both through difficulties in collecting observations but also since specimens are rarely collected, basic biological parameters-age/growth and maturity are generally unable to be established. In such situations, conservation listing advice must resort to use of data collected haphazard data collection or expert opinion (Friedman et al. 2018; Friedman et al. 2019). The limitations of this are clear and yet obviously remain difficult to overcome and hence typically remain in place as the best available basis for precautionary management. Indeed, elasmobranchs, rare or otherwise, exemplify this problem: Dulvy et al (2014) found that of 1041 species considered only 389 were not at risk of extinction. Crucially however, the study also noted that nearly half of all species were considered data deficient, clearly indicating a need for methods that can address severe uncertainty for a group which are broadly at risk due to their widely noted life history characteristics (Reynolds et al. 2005).

Close-kin mark-recapture (CKMR) (Bravington et al. 2016) is a technique that addresses many of these issues. It seeks to find closely-related kin-pairs in a sample of individuals from a given population. The number of close-relatives parent-offspring pairs or half siblings, is directly related to abundance of breeding-aged individuals in the population. CKMR has been applied to assess population abundance in a highly migratory marine teleost, *Thunnus maccoyii* (Bravington, Grewe, and Davies 2016) and for Australasian white shark, *Carcharodon carcharias* (Hillary et al. 2018; Bruce et al. 2018). A recent study (Ruzzante et al., 2019) has verified the approach against standard mark-recapture approaches. These studies have demonstrated the utility of CKMR in very different settings; a highly abundant, highly migratory teleost targeted by a multinational fishery; and a relatively scarce elasmobranch and a freshwater teleost. Kinship information has also been used to establish generational connectivity (Palsbøll, Zachariah Peery, and Berube 2010; Feutry et al. 2017)

In this study we examine an isolated population of elasmobranch, the speartooth shark (*Glyphis glyphis*) known to occur in only a few river systems throughout Northern Australia and Papua New Guinea (Pillans et al. 2009). Glyphis glyphis was described from a single specimen without locality and was subsequently found to be synonymous with the Bizant River shark (*Glyphis sp. A*). In Australia, the species was first recorded from the Bizant River in Queensland where it has not been detected since and is thought to be extinct. *Glyphis glyphis* is typical of many sharks and rays which use tropical estuaries and rivers for parts of their life cycle (Lyon et al. 2017).

The population of *G. glyphis* examined here occupies the Wenlock River system which lies on the western side of Cape York, Queensland and is subject to a highly seasonal flow regime (Leigh and Sheldon 2008). The system comprises the Wenlock and Ducie rivers and empties into the Gulf of Carpentaria (Figure 1). The genetic isolation of the local *G glyphis* populations was established by Feutry et al (2017) who used kinship data gathered from mitogenomes and genome-wide SNP data from the same samples considered in this study as well as samples from several rivers in the Northern Territory, far to the west of the Wenlock. Feutry et al., found strong support for river fidelity in juveniles in females. In these Northern territory rivers, males were found to father offspring in several rivers indicating a degree of movement. However, there were no kin-pair linkages from the NT populations and the Wenlock.

**Figure 1.**
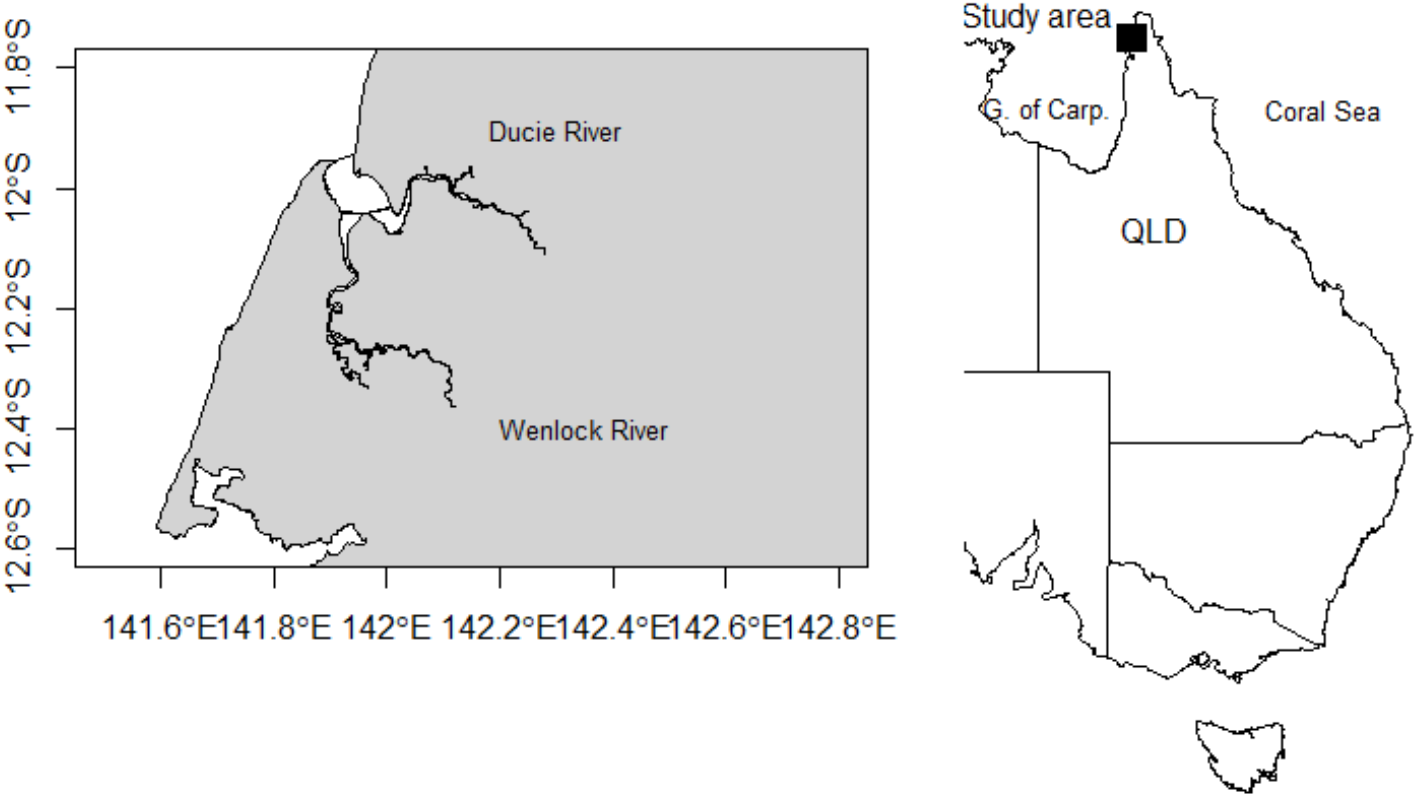
Map of Northern Australian region and (inset) details of the river system where samples were obtained. NT-Northern Territory and QLD - Queensland.

Like many euryhaline species that utilize estuaries and rivers as nursery areas, juvenile speartooth shark are confined to these habitats for several years before presumably moving into a marine environment. The six Australian and Papua New Guinea adults have only been recorded in marine environments (Pillans, unpublished data; White et al 2015). Members of the Genus attain large size (>2.5 m TL) and like other carcharhinid sharks of similar size and life history, most likely have low reproductive output and late maturation. However, since only six adults have ever been reported, virtually nothing is known about litter size and reproductive periodicity.

As we shall demonstrate in the following, CKMR based on half-sibling pairs detected within samples of mostly non-reproductive animals, produced estimates of adult abundance, sex ratio, and provide insight into reproductive periodicity and provide information on litter size from full-sibling pairs.

## Methods

### Sampling and genetics methods

Capture and sampling of *G. glyphis* was conducted under a CSIRO animal ethics permit (A11041; A2-2016; 2017-04). Juvenile *G. glyphis* (52-160 cm TL) were captured with rod and line in the estuarine reaches of the Wenlock and Ducie River (Figure 1). Neonate sharks were easily recognised by the presence of an umbilical scar and otherwise animals were assigned to an estimated age-class based on length and date of capture. We used a combination of growth data from neonate sharks recaptured over seven years and Given the rarity of the Data data on length at age were obtained from 10 G. glyphis from the Adelaide River, NT, that were aged from vertebral sections (P. Kyne, unpublished data).

The processing of the nuclear genetic data followed the method employed in Hillary et al (2018) and Bruce et al (2018), where the quality control, kin-finding statistical methods are outlined in detail. Briefly, this method compares every individual shark with all others which could feasibly be a related sibling and computes a Pseudo Log-odds (PLOD) score which characterizes likely kin-type as being either unrelated, full-sibling or half-siblings. Note that HSPs and grandparent-grandoffspring are genetically indistinguishable with unlinked SNP markers (Thompson 2000) and so age data needs to be employed to distinguish true HSPs. Mito-Genome sequencing was also examined using a PCR approach so that each HSP could be categorized as likely paternal or maternal.

### Estimates for adult abundance

The approach to estimating abundance and other population parameters proceeds by specifying the population dynamics, and then using maximum likelihood, to estimate the most likely abundance, survival rates etc. given the observed kin-pair data. In the following we describe the population model and outline the statstical estimation approach. Full details of the estimation methods are given in Appendix A.

### Population dynamics

The abundance dynamics were assumed to follow a simple exponential growth model split by sex:

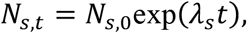

where *λ*_*s*_ is the logarithmic rate-of-change, *N*_*s*,0_ the initial abundance at some given reference year, and the annual adult survival probability is *ϕ*_*s*_. The subscripts *s* and *t* are sex and time, respectively. Note that we did not expect to see appreciable population growth over the series of cohorts captured by the data. Accordingly, and after initial model and data exploration, *λ* was set to 0, therefore we estimate *N*_0_ as a constant breeding abundance.

### Accounting for non-annual female spawning

It is known that many sharks skip-spawn - i.e. do not spawn annually - especially in females (Domeier and Nasby-Lucas 2007; Bansemer and Bennett 2009; Pillans et al. 2021). Note that, by skip spawning we do not mean a random chance that, in any given year, a female might choose to reproduce (or not) depending on individual factors such as body condition. That particular case does not bias the chance of finding HSPs (Bravington et al. 2016). Instead, we mean that the spawning cycle is fixed at a >1 year frequency for at least some proportion of the female population. For this systematic case, some fraction of the adult female population, *ψ* (maximum 1) operates on a non-annual breeding cycle. Here we allowed for a two year female breeding cycle. Therefore an unknown proportion of the population may spawn on even years and the remainder in odd. Assuming it to be a random process as to which cycle an individual belongs to, then of the total *N* adults, if we have *ψ* = 1 then only *N*/2 are on average ever reproducing in a given year. The effective adult reproducing population, 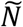, in a given year is a function of *ψ*.

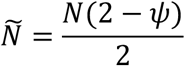

For *ψ* = 0 (every adult female reproduces every year), by definition 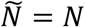.

The probability of being a maternal HSP now depends upon |*c*_1_ – *c*_2_|, where *c*_1_ and *c*_2_ are the cohorts in the comparison, so if |*c*_1_ – *c*_2_| is even, then following Hillary et al (2018), the probability of a maternal HSP (*MHSP*) is

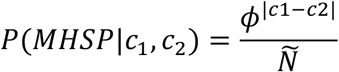

When |*c*_1_ – *c*_2_| is an odd number then

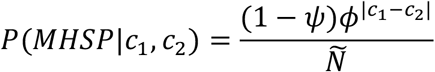

For the extreme case of population-wide skip-spawning (*ψ* = 1) there is zero chance of MHSPs for odd gaps between birth years and twice as much chance as finding HSPs for even numbered gaps because half the population is not reproducing that year. For the *ψ* = 0, then effectively *ψ* dissapears from the equations. In this study we examined this skip spawning model (spawning occurs every 2 years for proportion *ϕ* of females) and the non-skip model obtained by fixing *ψ* = 0.

### Abundance Log-likelihood

Estimating the log-likelihood of the parameters *λ, ϕ* and *N*_0_ requires comparison of each individual in the data to all others to determine if they are either an Unrelated Pair (UP), Full-sibling Pair (FSP) or a Half-Sibling Pair (HSP). In determining the probability of *i* and *j* sharing at least one parent, we must account for the survival rate of the adults over the time elapsed between their respective birth years. In doing this we also obtain information about the adult survival rate, *ϕ*^*A*^. Clearly, for an FSP to be found we expect that *i* and *j* are memebers of the same cohort, i.e. share the same birth year. Hence the key covariates for each {*i, j*} juvenile comparison are their respective years of birth: **z** = {*z*_*i*_, *z*_*j*_}. In brief the likelihood of the parameters is given by:

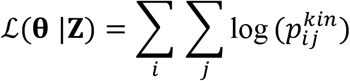

where 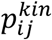 is the probability that individuals *i* and *j* are of a given kin-type and **θ** are unknown parameters to be estimated, including the abundance, mortality rate and population growth rate. Full details of the probabilities of kinship used in the likelihood calculations (see Appendix A).

In order to estimate abundance, growth or mortality in this model, various other factors of the sampling process that leads to kin pairs must be accounted for. This entails the following extra parameters in the model which in one sense are “nuisance” parameters, but in many respects also contain interesting information regarding the reproductive processes at play. Appendix A gives details of how these are factored into the likelihood calculations. Here we simply describe the rationale for their inclusion.

- Given that the model is structured by sex, we estimate the sex ratio *ζ*, the proportion of females in the adult population.
- There is high potential for over-representation of within-cohort maternal kin (FSPs and MHSPs) given high heterogeneity in very early-life survival. For example: some litters may by chance suffer zero mortality from predators, bycatch or other factors. Another may be subject to heavy mortality. We parameterise this through a term *v* which, following Hillary et al (2018), we term the “litter effect” parameter.
- Multiple paternities are likely given understanding of other elasmobranchs (Feldheim, Gruber, and Ashley 2004; Portnoy et al. 2007). The parameter *θ* is the probability that a mother will mate with more than one male in a given year.
- Similarly, *γ* is a parameter which describes multiple paternity. It is an estimate of the number of different females an individual father is expected to breed with in a given year.

Hence, the full set of unknown parameters is **θ** = {*N*_0_, *λ*, *ϕ*^*A*^, *v*, *γ*, *ζ*, *ψ*}. Estimation of these parameters was carried out using the R package ‘TMB’ (Kristensen et al. 2015). Posterior samples on model estimates were generated using the R package ‘tmbstan’ (Monnahan and Kristensen 2018).

Further, to consider the influence of sex ratio on the abundance estimates, we also employed a simple estimator of abundance accounting for sex ratio (see Appendix B) and used this to explore what effect misestimation of sex ratio (e.g. through uncertainty due to low mtDNA diversity) might have on adult abundance.

## Results

### Kin pair data

Data from 229 animals captured between 2013-2016 was used in this study. These consisted of 110 females and 109 males. The samples ranged between 48 and 226cm (Total length).

These samples generate *N*_*comp*_ = *N*_*samples*_ * (*N*_*samples*_ – 1)/2 comparisons between individuals and in this case, *N*_*comp*_ = 5995. From these the kin-finding methods based on the PLOD statistics (fig 2), that 5916 unrelated pairwise comparisons and the presence of 33 FSP and 46 HSP in the sampled individuals. Attributing age-at-length to these suggested that the earliest HSP was made up of an individual from the 2007 cohort. The cohorts observed in the HSPs spanned 6 years, with 76% being from cohorts within a 0-2 year interval. All full sibling pairs were from within the same cohorts (Figure 3). There were 23 kin clusters in the observed data and 15 FSP-only clusters. The maximum number of FSP’s from the same year was 5 individuals which were all litter mates. Given the probability of sampling all juveniles from a litter is low when only 30 samples taken in a year, it is possible that this provides an estimate of the lower bound on litter size for this species. The mtDNA data reflected very low haplotype diversity with only 2 haplotypes detected. These occurred in roughly equal proportions in all samples (type 1= 51.7%: type2 = 48.3%). Within FSP these were equally split again, but in HSP there was a preponderance of type 2 (type 1=37%: type 2 = 63%).

**Figure 2.**
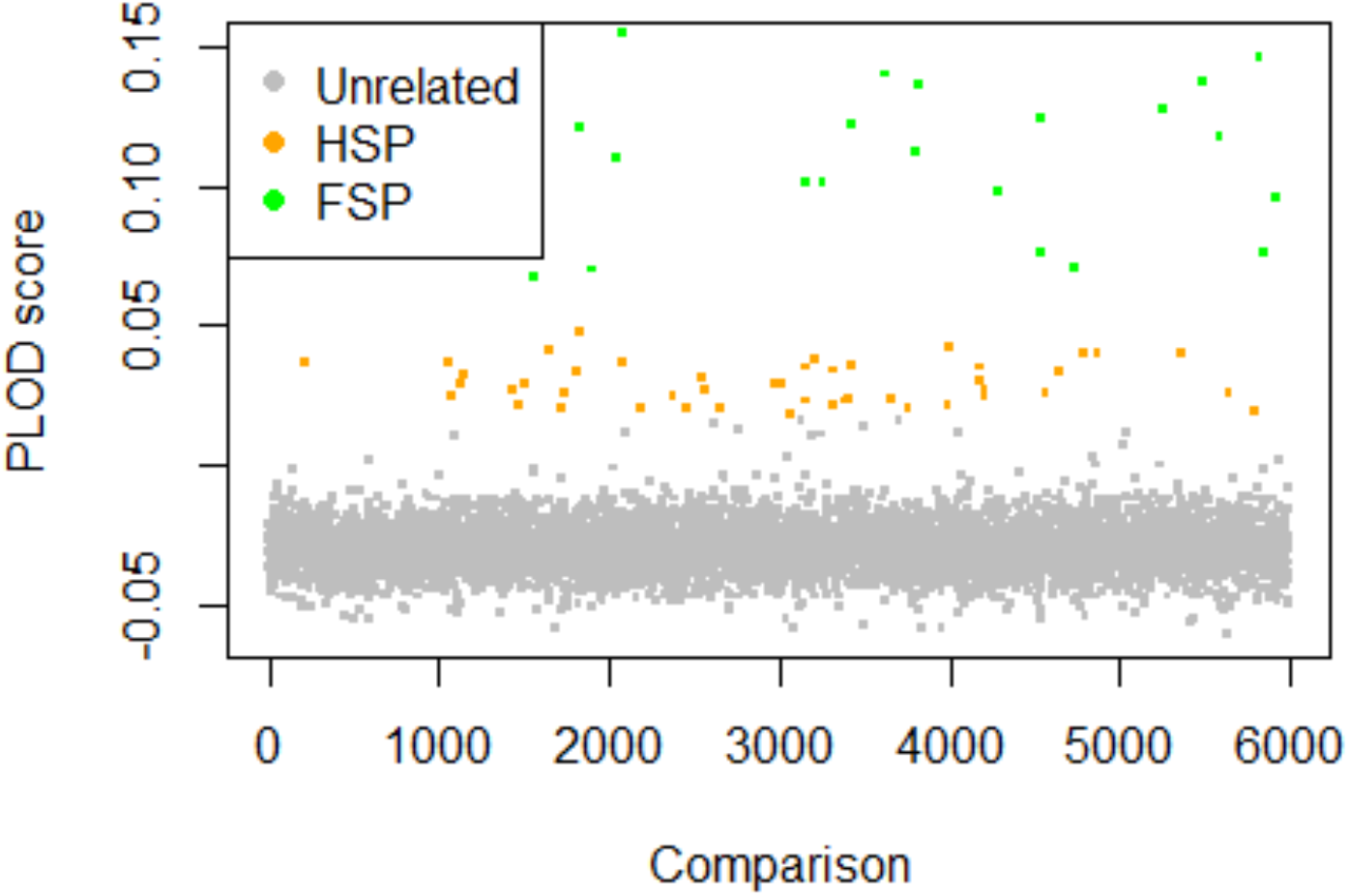
Psuedo-log odds (PLOD) scores for each pairwise comparison of the data. Colours denote kintypes.

**Figure 3.**
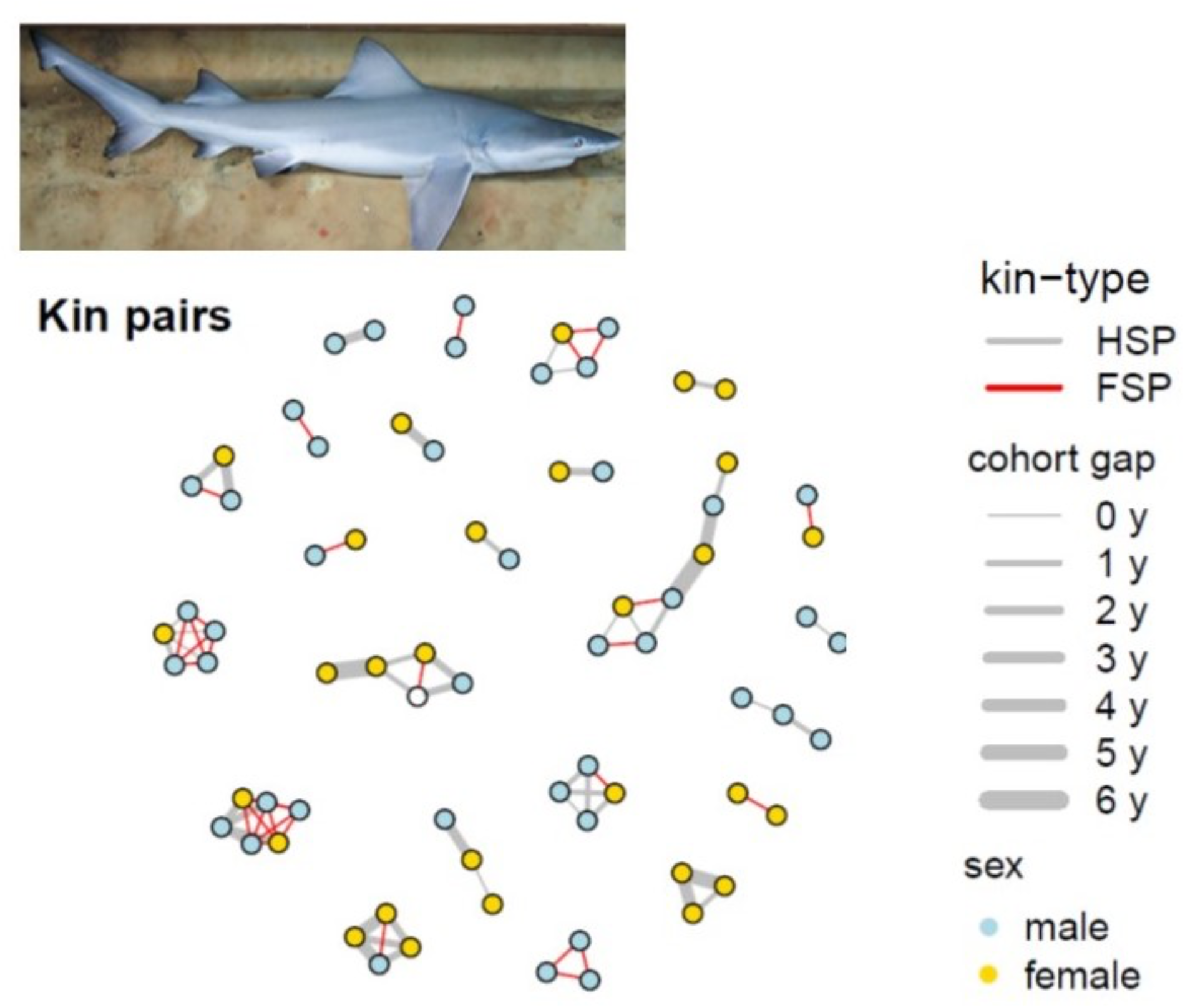
HSP and FSP Kin-relations detected from the 226 individuals. 23 clusters of related individuals were detected. Line weights denote the cohort gap in years between related individuals and the colours of edges denotes kin-type.

### Abundance estimates model estimates

The median-posterior estimate of adult abundance was 897 breeding individuals for the system with wide credible intervals (Table 1). The maximum a posteriori estimate at just over 500 (Figure 5a). We would therefore conclude the breeding population in the Wenlock river consists of 500-1000 individuals. However, the sex ratio parameter (Table 1) was highly skewed toward males with an estimated sex ratio of 92% male: 8% Female. Therefore the male abundance median estimate is 825.65 (95% CI: 358.67–2517.53), which imples that the number of breeding females is less than 100 individuals (Median: 72, 95% CI: 18-206). Again there is uncertainty around these estimates, but the model predicts a highly male–skewed sex ratio.

**Table 1.** Summary of model parameter estimates from posterior distributions. Values are given as median and 95% credible intervals.

| Parameter | Description | Median | 5% | 95% |
| --- | --- | --- | --- | --- |
| $N_0$ | Abundance | 897.45 | 365.99 | 3269.52 |
| $\nu$ | litter effect | 0.87 | 0.53 | 1.51 |
| $\zeta$ | sex ratio | 0.08 | 0.02 | 0.23 |
| $\theta$ | multiple paternity | 0.06 | 0.01 | 0.21 |
| $\gamma$ | female breeding partners | 3.83 | 1 | 16.93 |
| $\psi$ | skip spawning proportion | 0.7 | 0.41 | 0.85 |

To gauge the interaction between sex-ratio and abundance estimates, we also considered an alternate and simple approach using a more basic estimator of population size (see appendix B). This indicate similar results for estimated total adult abundance (≈ 850 adult individuals). Additionally we used it to determine what the likely effect of uncertainty or bias in our mtDNA results (specifically, the low diversity with only 2 haplotypes at 50:50 ratio). The results show that if the true sex ratio in the Wenlock is closer to 50:50, then we expect a smaller population of adults (at 50:50 sex ratio, the population would be roughly half what we estimate in the full model). This has important conservation implications which are considered in the discussion.

The multiple paternity parameter *θ* was low (0.06, 95% CI: 0.01–0.21). This indicates a low probability that breeding females mate with multiple partners in a given year. Accordingly the male-linked parameter *γ*, estimating the number of female partners per year for males was approximately 4 (95% CI: 1-17). The absence of cross-cohort FSPs indicates that there is apparently a low likelihood of individual mothers and fathers breeding together over different years. Additionally, the estimate of the skip spawning parameter strongly supported a large portion of the females obligately skip spawning. The skip spawning parameter was estimated at *ψ* = 0.7 (95% CI: 0.41 – 0.85). Note that cross cohort HSP at a 1 year gap were most common (1-2 year gaps accounted for 52% of the observed HSPs).

### Model fit diagnostics

Two diagnostics were used to assess model fit. The first used the HSPs and haplotype data to consider the predicted and observed numbers of HSP within and across cohorts that had the same or differing haplotypes. There was very good agreement between the model predictions and the observed numbers (table 2). The second diagnostic looked that the expected number of HSPs that would be expected for a given cohort gap (Figure 4) which again showed close agreement between the model predictions and the observed numbers of HSPs at a given number of gap years between cohorts. From these results we consider that the model fits the data well giving confidence in its predictions.

**Table 2.** Diagnostics of expected vs observed numbers of HSPs that did or did not have a common haplotype. The model predictions agreed closely with the data.

| Cohort | Within |  | Across |  |
| --- | --- | --- | --- | --- |
| Haplotype | Same | Different | Same | Different |
| Observed | 40 | 4 | 33 | 2 |
| Predicted | 40.001 | 3.999 | 32.986 | 1.999 |

**Figure 4.**
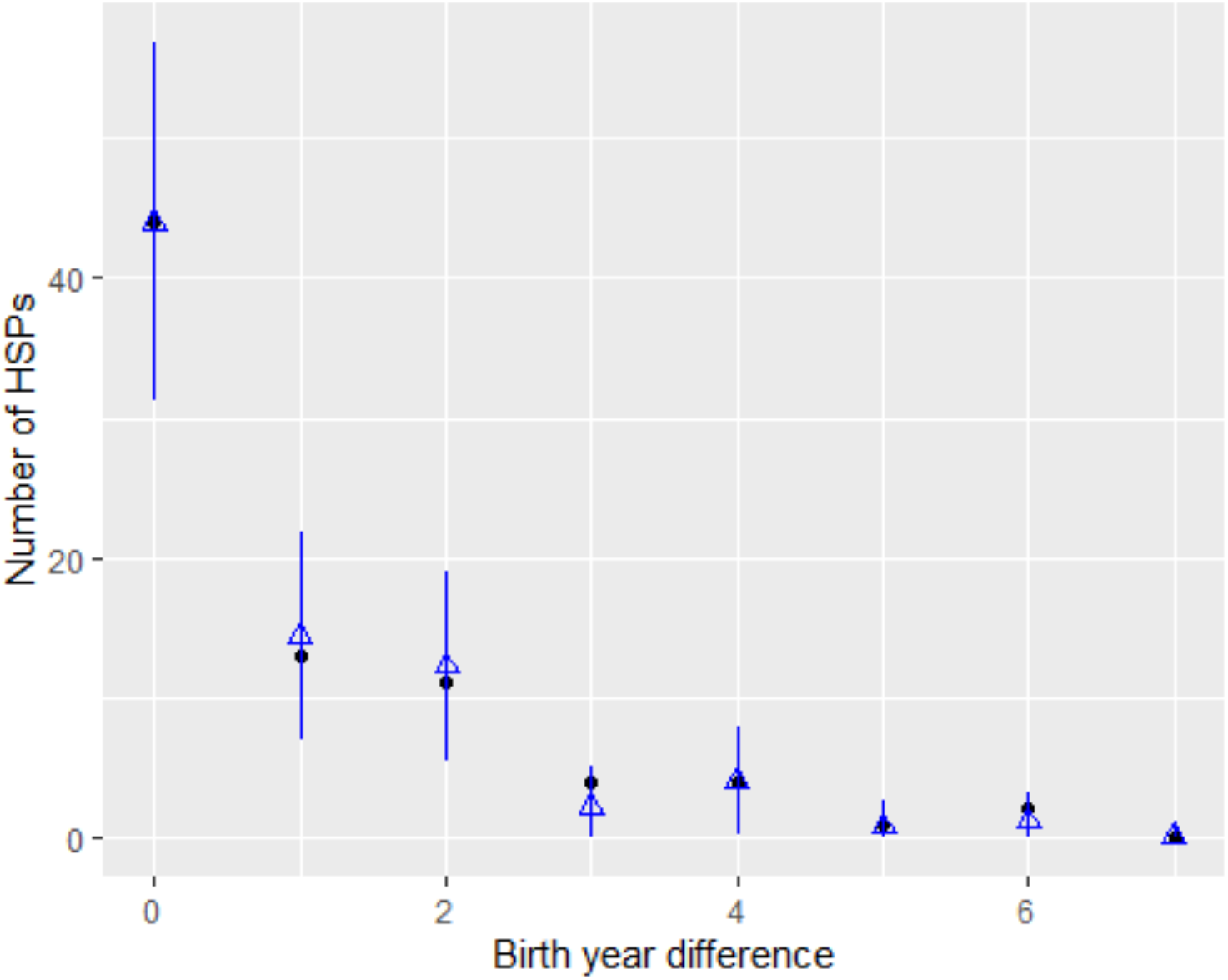
Diagnostics of model fit. The observed number of HSPs are given by black dots and expected numbers are given as blue triangles plus standard errors

**Figure 5.**
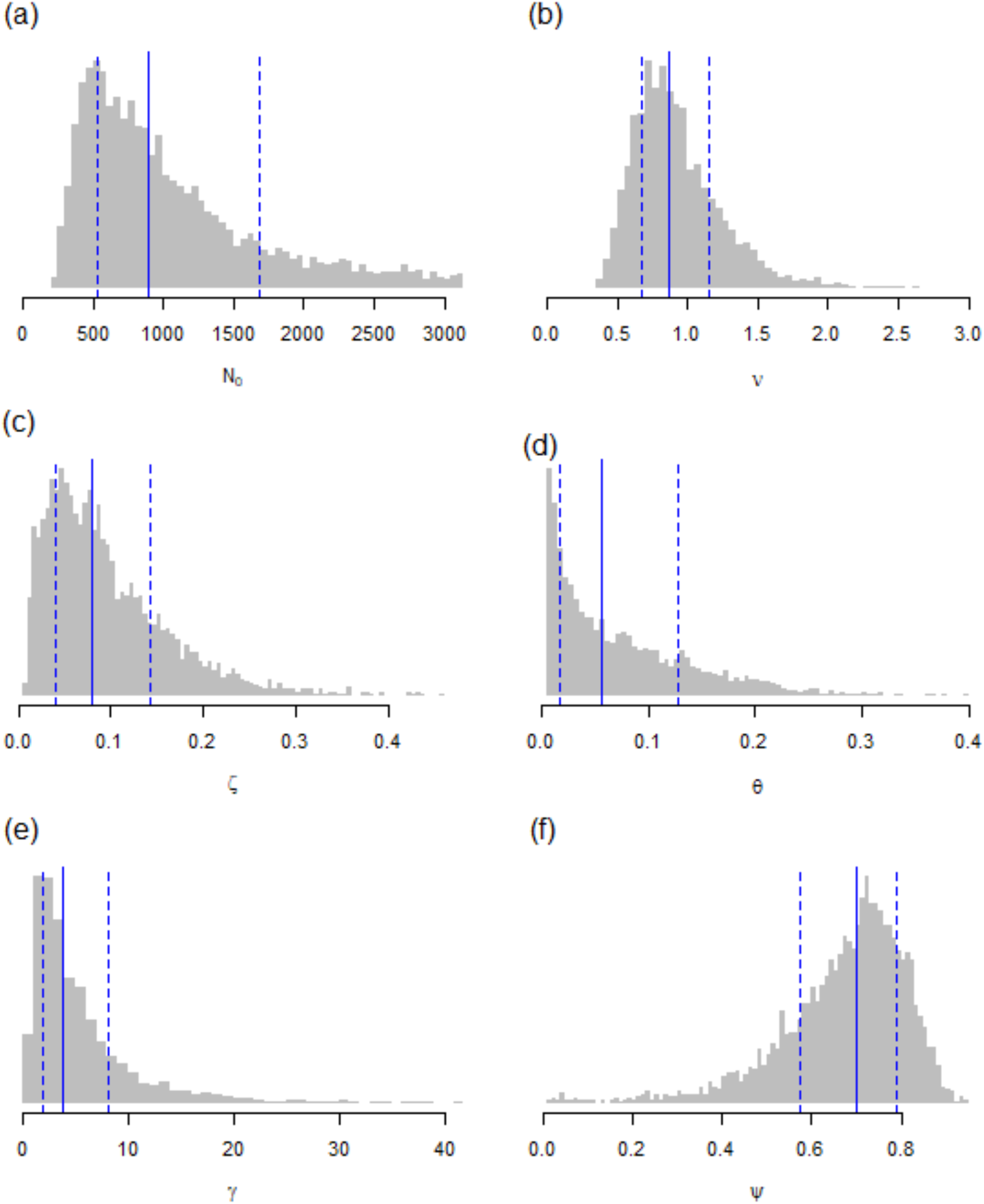
Posteriors on model estimates (a) abundance *N*_0_ (b) The litter effect *v* (c) The sex ratio *ζ* (d) *θ* multiple paternity (e) female partners per female estimate *γ* (f) proportion of females skip spawning *ψ*. Vertical bars indicate median (solid line) and 80% credible intervals (for 95% CI summaries, see table 2).

## Discussion

Elasmobranchs are ackowledged to be a taxa at high risk of population decline through harvesting and habitat loss (Dulvy et al. 2014). This situation is exacerbated by uncertainty with more than half the species assessed classified as data deficient (Collen et al. 2016). Reliable calculation of either relative or absolute abundance is problematic for a great many species. River sharks, prior to this study could be considered to be an extreme exemplar of these difficulties. However, we showed through this study that CKMR is able to produce a robust estimate of abundance of breeding individuals and other parameters of interest over the course of a 4 year sampling program.

Maintaining a viable breeding population is obviously a crucial conservation target, yet it is not an uncommon situation in elasmobranchs and other species, that assessing the number of breeding adults via direct marking or survey methods is impossible. Our result also demonstrates that it is possible to rigorously assess endangered and elasmobranchs where the adults are effectively cryptic, so long as juvenile samples are available. That field programs were able to successfully capture these individuals and thereby estimate the number of breeding individuals means that these techniques could be applied to other threatened species.

Threats to *G glyphis* neonates in the Wenlock include natural local predators, such as the saltwater crocodile (*Crocodylus porosus*) and bull sharks (*Carcharinus leucas*) and fishing pressure from gill netting and crab trap-fishing (Pillans et al. 2009; Lyon et al. 2017). It is known that *G glyphis* are captured in these gear types (Pillans et al., in press) and typically tend to be mistaken for small *C leucas*. Given the low breeding female abundance from the present study, there is strong potential for juvenile take to have adverse effects on the population in coming years. This is clearly an aspect of the management of the population which requires further research and monitoring. We recommend studies that investigate juvenile mortality, juvenile abundance and the risk posed by bycatch in commercial gill nets and commercial and recreational crab fisheries that operate in the system.

A key result from our study that is the highly male-biased sex ratio we estimated based on the mitochondrial haplotypes in FSP and HSP. There is considerable uncertainty around this result, and further data is required to refine our estimate of sex ratio and the prevalance of skip spawning. Preliminary work where we set the skip-spawning parameter *ψ* = 0, indicated that the overall adult abundance estimates remained largely unchanged, whether skip spawning was included or not. The effect of including skip spawning was to increase the degree of male bias in the sex ratio as skip spawning by females would reduce the average number of females breeding in any given year.

Given these results, and the potential for the estimation process to trade-off increased precision on one parameter at the expense of another, we critically examined the skewed sex ratio result by considering some simple facts from our data. Taking the across-cohort case only, the two key data were as follows: (1) Two haplotypes with exact 50/50 frequency in the population (which, while unusual, makes the mathematics simple), and (2) 33 of the cross-cohort HSPs the same haplotype while 2 did not. Consider a simple case with a stable total adult population *N*_*A*_ and proportion of females *ζ*. Mortality can be ignored if we assume it is equal for each sex. Then in this simple case, the probability of being a maternal HSP is then 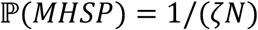 and paternal 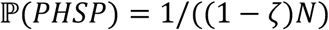. Now consider how we might relate this to mtDNA data.

When comparing two HSPs with haplotypes *h*_1_ and *h*_2_ (in our case, the only ones) then the probability that they do have the same haplotype is:

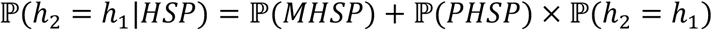

The probability they have different haplotypes is:

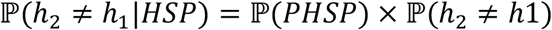

From this one can examine what would constitute a 50/50 sex ratio in the adults by setting 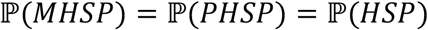. Knowing that 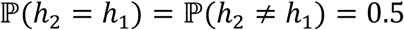 from the 50:50 haplotype frequency in the population, then the probability (assuming 50/50 sex ratio) of getting a HSP that either (a) has a common haplotype is

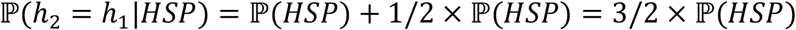

or, (b) has different haplotype is

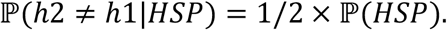

The cases (a) and (b) imply that even **with** a 50/50 sex ratio, the expected ratio of cases that do/do not have the same haplotype is already 3:1. If there are effectively an infinite number of haplotypes in the population, then a 50/50 sex ratio will converge to a 1:1 ratio in numbers of HSPs that do or do not have the same haplotype. But when the haplotypes become rarer, this ratio immediately starts to rise from 1:1 to much higher values. With only 2 haplotypes each having 50% frequency in the population, we see a ≈ 16:1 ratio in the number that do/do not have the same haplotype, indicating far fewer females than males. Which is as we see in our model estimates.

In fact, we can produce a simple estimate of sex ratio, purely from the observed mtDNA data (i.e. with 2 haplotypes at 50% frequency). The ratio of HSPs that have the same haplotype to those that do not is (2 – *ζ*)/*ζ* so *ζ* = 0.5 gives a ratio of 3. If we let *χ* be our observed HSP ratio, then our guess at the sex ratio is *ζ* = 2/(1 + *χ*) so substituting in *χ* = 16 we obtain *ζ* = 0.11 which was our model estimates if we set the skip-spawning proption to zero. This alternative argument supports the finding that the sex ratio of *G. glyphis* in the Wenlock is highly skewed, given the data on proportion of the HSP which had common haplotypes and the fact that only 2 mtDNA haplotypes were observed.

In our results there were two key findings regarding breeding biology:

First, we estimated a distinctly male bias via the sex ratio parameter *ζ*. But *θ*, the female probability of multiple male partners is estimated to be very low *θ* = 0.06 (95% CI: 0.01-0.21), while *γ*, the number of female partners per male is ~4 (95% CI: 1-17). The skip spawning proportion *ψ* ≈ 0.7 and therefore, on average, the model predicts that there are ~45 breeding females in the system for a given year.

Second, if mating was totally random then in the absence of any other information besides a sex ratio and abundance, we would expect that females might mate with multiple males (the estimates at face value suggest there are ≈ 11.5 males per female in the population). However, the extra information from the kin pairs and mtDNA in the model implies that this was not the case. In fact these data suggest that female promiscuity is rare whereas males mate with multiple females. But also given magnitude of *γ* and the estimates population size, the model indicates that not all breeding males sire offspring in a given year.

A situation where females mate with few partners and then do not mate again within that season would be consistent with these findings. Note that the litter effect parameter is not overly large (*v* = 0.8-0.9), therefore indicating that there is not an overwhelming effect of “lucky litters.” Given that, we can use the estimated sex ratio and number of female breeding partners per year (*γ*) to calculate the effective/expected number of breeding males as 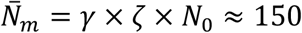. This would equate to a “within year” sex ratio of 22% F: 78% M. Clearly this is still skewed toward males, but not to the extent that simplistic interpretation of the parameter estimates would indicate.

Male-biased sex ratios do occur in vertebrate populations and examples exist in birds, mammals and reptiles (Clutton-Brock 1986; Cockburn et al. 2002), although in most populations, the Fisherian ratio is expected (Hamilton 1967; Charnov 1982). Occurrences of sex ratio bias have been implicated in increasing risk to endangered populations (Heinsohn et al. 2019). In certain reptiles, male aggression during courtship has been noted to have the effect of reducing female survival rates (Le Galliard et al. 2005). In turn, the evolutionary response by breeding females was to produce more male offspring as they would be expected to have higher fitness. This could be an explanation for our results, given that sexual conflict has been observed in elasmobranchs (Portnoy et al. 2007). However, since adults are virtually never seen, the potential to observe or test such a mechanism is almost impossible in this case.

The estimates obtained here are consistent with a roving population of males and philopatric females. Female philopatry has been observed in other Australian *G. glyphis* populations (Feutry et al 2017) and more widely in elasmobranchs (Hueter et al. 2005; Portnoy et al. 2015; Feldheim et al. 2014; Feldheim, Gruber, and Ashley 2004; Nosal et al. 2021). Thus female philopatry would be expected in the Wenlock. However, a wide ranging adult male population would be expected to breed in other river systems which would be expected to show up as between-river kin pairs. Feutry et al (2017) found no kin-pairs were between the Wenlock and other known Australian populations. The data from Feutry et al (2017) did not include samples from Papua New Guinea and it is possible other rivers around the Gulf of Carpentaria which may hold *G. glyphis* and that the Wenlock population is actually reproductively connected to other locations. Hence, these results indicate a need for further sampling in other systems. In the Northern territory *G. glyphis* populations, males sire offspring between rivers but females were philopatric to the same river each year (Feutry et al., 2017). Such a system might explain the male-skewed sex ratios in this study, but again there are no suitable rivers north of the Wenlock apart from those in Papua New Guinea. We consider this unlikely given the previous results from the Northern Territory where populations 150 km shared a parent – but not in the same year. South of the Wenlock the nearest large river is over 160 km away with no records of juveniles observed in this system. Additionally, if unknown populations of *G. glyphis* were connected to the Wenlock population, it would most likely be via males with females being philopatric, as per the results of Feutry et al (2017). Therefore female recruits would only originate in the Wenlock itself.

These aspects mean at most, there could be some unknown connectivity of the Wenlock population by males only. Therefore reduction of numbers of females in the system would not be offset by immigration. This suggests that the area is critical habitat which needs careful management and monitoring on an ongoing basis, in common with many elasmobranch nursery areas (Heupel et al. 2019). Previous studies have considered detailed spatial management schemes for this population (Dwyer et al. 2019). However, these schemes require detailed fisheries effort data and associated bycatch rates. Bycaught *G. glpyhis* are likely to be largely the same juveniles and sub-adult age classes sampled here. Therefore, further investigation of juvenile population abundance and mortality will be needed in order to predict long term impacts on the breeding stock and the sustainability of the population.

## Conclusions

This study has provided a robust estimate of the breeding population of an extremely poorly understood and rare elasmobranch. Our model provided insights into the breeding dynamics of the population and produced some surprising results regarding the sex ratio in this population. Clearly, these will require further investigation which may revise the current estimates (Cockburn et al. 2002). Nonetheless, our abundance estimates were robust to these aspects and point to the need to obtain information on sources of juvenile mortality in the system as well as the potential for linkages to other populations. At the moment, any such conclusions regarding another additional Queensland population would be premature and the proper precautionary approach would be to continue to consider the Wenlock as an isolated population with as few as 100 adult females. This and the potential for significant mortality on juveniles from fisheries recently found in Pillans et al (in press) indicate a need for ongoing protection and monitoring.

## Acknowledgements

This work was funded by the Ord River Offset project and the CSIRO.

## Author contributions

RP conceived the study, collected field samples and conducted tagging work; PF, RG, JM performed laboratory genetics work; TAP, RH conducted analysis and statistical modelling and wrote the manuscript. All authors reviewed the manuscript prior to submission.

## Supplementary Material

## Appendix A CKMR likelihood calculations

To define the various combinations of HSP probabilities we first consider the within-cohort (*c*_*i*_ = *c*_*j*_) and then consider cross cohort situation. Throughout we define where the set of kintypes as 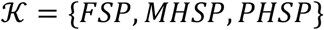. The following terms are used in the likelihood.

- The *v* term is the litter effect. This describes the potential for over-representation of within-cohort maternal kin (FSPs and MHSPs) given high heterogeneity in very early-life survival. For example: some litters are by chance missed by all predators such as crocodiles, whereas another may be subject to 100% predation.
- The *θ* term is the probability that a mother will mate with more than one male in a given year i.e. multiple paternity litters.
- The *γ* term is present to permit the possibility of individual fathers mating with multiple females in a given year
- The HSP false-neg probability 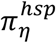 is included to take care of the fact that we know we removed a certain fraction of true HSPs away to ensure we avoid false positives in the final sample.
- The critical HSP PLOD *η* is the threshold such that we expect less than 1 non-HSP appearing above it; for this example 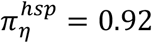 (with HTPs being the critical kin for defining where *η* was). For FSPs there it was clear from their PLODs scores relative to other kintypes so we assumed that *π*^*fsp*^ = 1.

We now consider the cases of within and cross-cohort comparisons and their associated kin-pair probabilities.

## Within-cohort comparisons

For the Full-Sibling Pair (FSP) case:

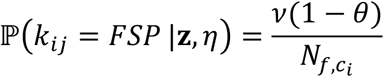

and for Maternal HSPs (MHSPs):

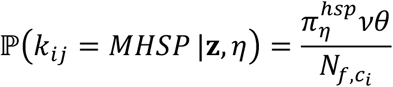

Now consider Paternal HSPs (PHSPs):

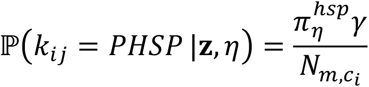

## Cross-cohort comparisons

Dealing with the cross-cohort case now (*c*_*i*_ ≠ *c*_*j*_) then for MHSPs (assuming cross-cohort FSPs to be basically zero probability):

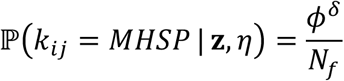

where *δ* = *c*_*i*_ – *c*_*j*_. The same general expression applies to cross-cohort PHSPs (just swap *f* for *m*). The overall negative log likelihood is therefore given by

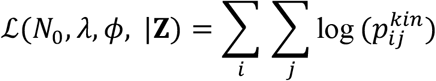

where

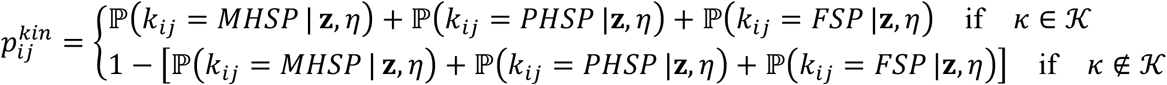

## Inclusion of mitochondrial DNA

Mitochondrial DNA provides information on the following question: given the covariates, **z**, and the critical PLOD, *η*, do the compared individuals have the same haplotype or not?

The approach we take is to split the cases into three groups:

- Case 1: Probability that you are above some PLOD value *η*′, where *η*′ > *η*, and are a FSP only (and therefore both must have the same haplotype, i.e. *h*_2_ = *h*_1_)
- Case 2: Probability that the PLOD is between *η* and *η*′ (i.e. and are HSP) and *h*_2_ = *h*_1_
- Case 3: Probability that your PLOD is between *η* and *η*′ (i.e. and are HSP) and the haplotypes differ (*h*_2_ = *h*_1_)

The are two steps to populating the probabilities that go along with these three cases: (i) calculate them as ‘raw’ probabilities given the PLOD and covariates for each comparison above the critical PLOD, *η*; and (ii) normalise them to form a categorical distribution with the three cases as outlined above. For case 1 (PLOD > *η*′) the raw probability is just 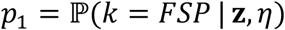. For case 2 (PLOD ∈ (*η*, *η*′)and *h*_2_ = *h*_1_) is is defined as follows:

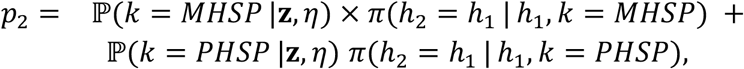

and for case 3 we have that

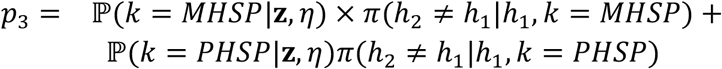

For MHSPs *π*(*h*_2_ = *h*_1_ | *h*_1_, *k*) = 1 and *π*(*h*_2_ ≠ *h*_1_ | *h*_1_, *k*) = 0; for PHSPs *π*(*h*_2_ = *h*_1_ | *h*_1_, *k*) = 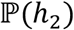 and 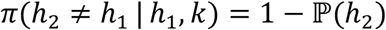. These three probabilities are then normalised to sum to 1

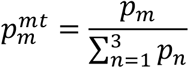

So for every case, *m*, where we have an identified kin-pair above the critical PLOD *η* we add 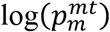 to the log-likelihood, depending on which of three cases happens to cover that particular comparison that found the kin-pair.The last bit of the likelihood (which is Bernoulli) will then be the nuclear DNA component describing the probability of being “above the *η* line”:

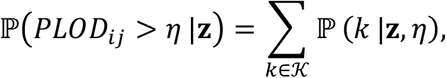

computed over all possible {*i, j*} comparisons.

## Appendix B. Simple estimator of abundance from CKMR accounting for sex ratio

First, note that we must use only across-cohort comparisons 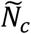 and HSPs 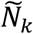 to avoid biasing abundance calculations. However in the present case we need to account for the sex ratio *ζ* given we have indications of a significant skew. Following Hillary et al (2018), the further components we need (and the values from our study) are: 1. Mean difference in birth years of across-cohort comps *δ* (2) 2. Adult survival rate (assumed same across sexes), *ϕ* (0.8807971) 3. False negative retention probability, *π*_*η*_ (0.92) We also set *ζ* = 0.11 and note that we have *N*_*c*_=4094 and *N*_*k*_= 35. From these quantitites, the approximate probability of getting the total number of HSPs is as follows:

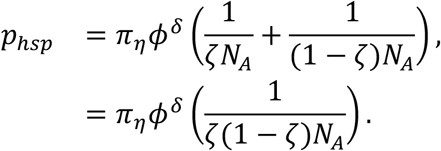

Recall that there is a simple relationship between the number of HSP and the number of comparisons:

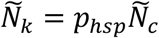

Now we use this to get an estimate of *N*_*A*_:

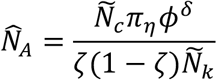

So in the Fisherian case, *ζ* = 0.5 and the equation contains the standard factor of 4;

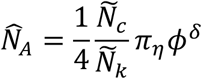

as per Hillary et al (2018). These calculations give an estimate of abundance as 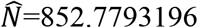, which is close to our estimate from the full model (see table 2 and figure 4 in the main text). Note also that as the sex ratio becomes skewed toward dominance by either sex, the population size goes up and that the lowest population size estimate is at a 1:1 sex ratio.

**Figure S3.**
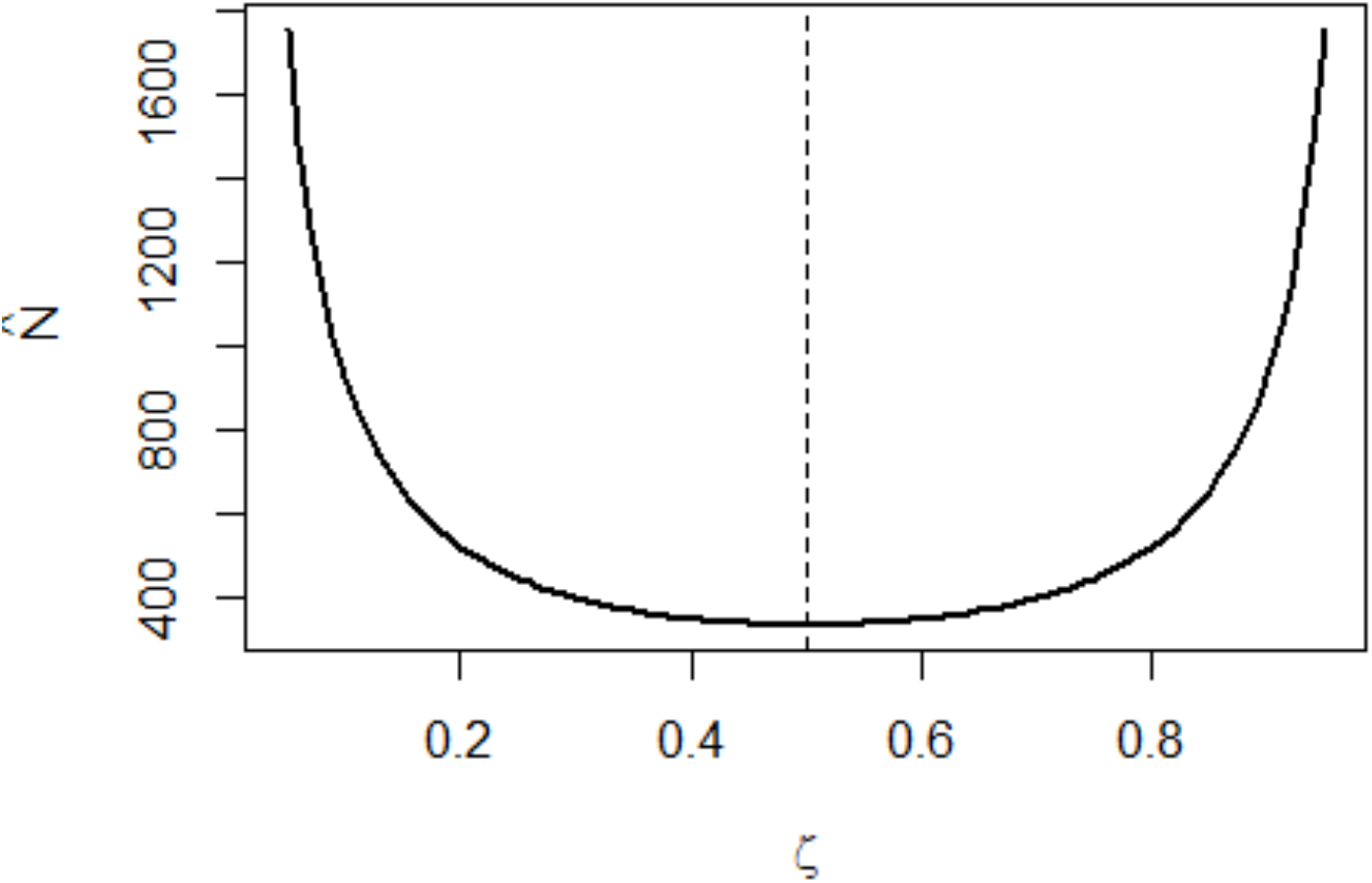
The relationship between sex ratio *ζ* and the estimated population size 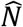 from the simple estimator and with values derived from our data. This shows that the lowest population size would be expected at 50:50 sex ratio.

